# Neural Dynamics of Tonic Cold Pain: A Novel Investigation of an In-Scanner Alternative to the Cold Pressor Test in Healthy Individuals

**DOI:** 10.1101/2024.12.03.626599

**Authors:** Sonia Medina, Sam W. Hughes

**Affiliations:** Department of Clinical and Biomedical Sciences, Faculty of Health and Life Sciences, University of Exeter

**Keywords:** Pain, cold pressor task, fMRI, ICA, seed-based, DCM

## Abstract

The cold pressor task (CPT) is widely used to study tonic pain during acute and chronic conditions and is often as used as a conditioning stimulus to activate descending pain control systems. However, logistical challenges in magnetic resonance imaging (MRI) limit its application, hindering the understanding of CPT’s neural dynamics. To address this, we acquired resting-state functional MRI data from 30 healthy participants before, during, and after immersion in gelled-cold water, the closest in-scanner alternative to date to CPT for prolonged stimulation. Participants provided subjective pain intensity ratings after each scan, as well as average pain perceived during noxious stimulation, using a numeric rating scale (NRS). Following fMRI, participants rated their pain continuously during identical tonic noxious stimulation of the contralateral hand using a visual analogue scale (VAS). We employed three complementary methods to examine changes in brain function across fMRI conditions: a data-driven approach via independent component analysis (ICA), seed-to-whole-brain connectivity analysis with the periaqueductal grey (PAG) as seed, and spectral dynamic causal modelling (spDCM) to explore effective connectivity changes across the dorsal anterior cingulate cortex (dACC), anterior insulae (AI), thalamus, and PAG. NRS scores were significantly higher following tonic cold compared to baseline and recovery conditions. Continuous VAS reflected sustained mild-to-moderate pain over six minutes, with average VAS scores not significantly differing from NRS ratings recorded in the scanner. ICA identified engagement of descending pain control and sensorimotor networks during pain, with the latter persisting during recovery. Seed-based analysis revealed a disengagement between the PAG and cortical/subcortical regions involved in pain processing, such as the dACC, midcingulate cortex, AI, intraparietal sulcus, and precuneus. Finally, spDCM revealed tonic pain neural signature was most likely characterised by top-down inhibitory and bottom-up excitatory connections. This study establishes the cold gelled-water paradigm as a robust in-scanner alternative to CPT. By uncovering key neural dynamics of CPT, we provide new insights into the brain and brainstem mechanisms of tonic cold pain paradigms routinely used in psychophysical pain studies.

**Key points:**

- Immersion in gelled-cold water is a reliable in-scanner substitute for the cold pressor task, enabling prolonged tonic cold pain research in MRI.
- Key neural dynamics, such as PAG-driven excitatory inputs and AI-mediated inhibitory control, were identified, providing new insights into cold pain modulation.
- By taking a multi-method approach, including ICA, seed-based connectivity, and DCM, we offer a comprehensive view of the neural networks involved in tonic cold pain, bridging neuroimaging and behavioural research.

## Introduction

The cold pressor task (CPT) is an experimental approach used to induce tonic pain in healthy participants [17; 20; 28; 63; 77; 81] and chronic pain patients [32; 33; 51; 53; 58; 62; 72; 75]. The responses to CPT are typically measured psychophysically, producing mild – moderate pain intensity and unpleasantness ratings [19]. CPT is also routinely used as the most common conditioning stimulus to activate key descending pain modulatory pathways as part of conditioned pain modulation (CPM) paradigms [34]. These CPT responses are therefore likely associated with enhanced bottom-up and top-down nociceptive signalling that shape both the pain experience and adaptive response to pain. Thus, detailed assessment of the neural dynamics underlying these responses is essential to decipher how tonic cold pain shapes complex cortical, sub-cortical and brainstem pain processing pathways.

Functional magnetic resonance imaging (fMRI) represents a useful tool to examine the neural underpinnings of tonic noxious stimulation [42]. Through measurements of blood oxygen level-dependent (BOLD) signals, fMRI captures local changes in brain activity during a task, but also allows for dynamic tracking of BOLD fluctuations across prolonged conditions such as tonic pain, an approach called resting state fMRI (rs-fMRI) [69]. Critically, replicating CPT-like experiments in an MRI environment poses considerable logistical and safety challenges; immersing a hand in water within the scanner bore introduces risks associated with placing conductive fluids near the powerful MRI magnetic field, potentially compromising both safety and data quality [66]. Consequently, a handful of fMRI studies have used alternative cold stimuli, such as gel pads [25] or aluminium cuffs [27; 43], targeted different body sites like the foot [25; 30; 61], carried out cold noxious stimulation separately from fMRI assessments [13; 24] or employed block designs with short lasting periods of hand immersion [36]. These efforts have undoubtedly advanced our understanding of the neural mechanisms underlying tonic cold pain in healthy population, pointing towards a disruption of descending pain control and autonomic control pathways via modulation of activity in the periaqueductal grey (PAG) [27; 36; 43], amygdala [13; 24; 61], insular cortex and basal ganglia [35]. Nevertheless, experimental setups that deviate from traditional CPT paradigms inevitably limit the generalisability of the results, hampering our ability to understand brain mechanisms underlying results from the psychophysics literature. Furthermore, the methodological choices to analyse fMRI data across the aforementioned studies varied greatly, ranging from block designs to rs-fMRI, from whole brain exploratory results to seed-based analyses constrained to a few regions of interest, from BOLD fMRI to arterial spin labelling fMRI, from 3 to 7 tesla MRI scanners. This absence of methodological and analytical consistency further impacts the comparability, generalisability and interpretability of the results.

Lapotka et al. [37] developed an MRI-compatible alternative to the cold pressor test by adding a thickening agent to the water, creating a gel-like substance. This method, used to successfully induce experimental pain in women with chronic pelvic pain, demonstrated BOLD activation in pain-processing brain areas within an evoked-responses fMRI block design. However, the impact of prolonged tonic cold noxious stimulation through gelled-cold water hand immersion on brain dynamics during rest in healthy participants remains unexplored.

In this study, we examined the effects of hand immersion in noxious gelled-cold water on rs-fMRI dynamics in 30 healthy participants, measuring BOLD responses during six minutes at baseline, during tonic pain, and in recovery. We employed three complementary data analysis methods: a data-driven, fully exploratory independent component analysis (ICA), a seed-based connectivity analysis, and a model-driven approach using spectral dynamic causal modelling (spDCM). We proposed that this multi-method approach could allow us to maximise interpretability by integrating insights from different analytical perspectives. Through this strategy, we not only provide a robust account of the neural dynamics associated with tonic cold pain, but also offer a framework for future studies to understand how methodological choices impact the interpretation of rs-fMRI data in pain research.

## Materials & Methods

### Experimental subjects

Thirty healthy, pain-free participants were recruited for this study (18 female; mean age= 29 years, SD=10). One participant was excluded from data analysis for not completing the MRI session due to high anxiety during scanning. All participants had no history of neurological or psychiatric conditions, no history of acute migraines or concussion resulting in loss of consciousness and no history if dizziness or motion sickness. Additional exclusion criteria included history off substance or alcohol abuse, inability to lie still within the MRI scanner environment, presence of medication that may affect temperature sensitivity, endogenous analgesia or cardiovascular medications, inability to understand instructions in English, and presence of any additional MRI contraindication (e.g., presence of metal in body, pacemaker, pregnancy, etc). 24 hours prior to their visit, participants were instructed to abstain from alcohol or any recreational drugs, to abstain from painkillers or paracetamol for 12 hours before their visit and to limit themselves to a maximum of one caffeinated drink in the morning. At the beginning of each visit, participants were asked to confirm adherence to these guidelines. If any guidelines were not followed, the visit was immediately terminated and rescheduled. Participants provided written informed consent at a different visit. The experiment was approved by the Health Research Authority and Health and Care Research Wales ethics committee (Ethics reference: 22/HRA/4672). Participants were given the opportunity to withdraw from the study at any point.

### Procedure

Participants attended five sessions in total (sessions 1-4 reported elsewhere) and one single MRI session. MRI scanning took place at the Mireille Gillings Neuroimaging Centre facility in Exeter. At the beginning of the MRI session, compliance with lifestyle guidelines and MRI eligibility were assessed. Participants also completed the state part of the State Trait Anxiety Inventory [70] to assess participants anxiety levels on the day. Next, participants underwent localiser and structural scan, followed by three resting state functional MRI (rs-fMRI) scans, each lasting approximately six minutes. The first rs-fMRI scan was a *baseline*, just aimed to assess markers of brain function at rest in the absence of additional stimulation. The second scan represented the *tonic* pain condition; before the scan began, a cold gel tub was placed on the participants’ stomach over a plastic sheet, in order to avoid cooling participants’ skin. Participants were instructed to fully submerge their right hand in the gel as soon as scanning began and keep it still inside the gel for the duration of the scan. Once the scan terminated, participants were instructed to take their hand out of the tub, and the experimenter entered the scanner room to remove the tub. The final scan mimicked the baseline scan, and was aimed to assess brain function during tonic pain *recovery.* At the beginning of the baseline scan and at end of each scan, participants answered the question ‘in how much pain are you right now?’ with a numerical rating from 0 (no pain) to 100 (worst pain imaginable). In addition, following the *tonic pain* scan, participants reported their perceived average pain during tonic cold stimulation with numerical rating from 0-100. During each rs-fMRI scan, participants were instructed to lie still with their eyes open and focus on a white fixation cross on a black background displayed at the centre of the screening in front of them. To further understand the temporal dynamics of participants tonic pain throughout the *tonic* pain condition, participants were presented with the same tonic cold stimulation following fMRI on their left hand, in order to avoid carry over effects on their right hand, and they were instructed to provide continuous subjective pain ratings across 6 minutes on a computerised visual analogue scale (VAS) [9] anchored with ‘no pain’ and ‘worst pain imaginable’).

### Materials and Measures

#### Cold gel

The method to deliver noxious tonic cold stimulation followed the strategy implemented by Lapotka et al. [37]. The gel was a mixture of 300g of cornstarch, 2 litres of water and 100g of salt, mixed in a 2-litre standard, rectangular plastic food container. A pilot testing demonstrated that the temperatures employed by Laptoka et al for testing (<1°C) were not bearable for most participants included in pilot testing for the full duration of the scan, therefore the tubs were kept in a fridge set to 5°C for at least 24 hours before each scanning visit, allowing for tonic stimulation that was bearable for six minutes while eliciting different levels of tonic pain. In order to keep the initial temperature as constant as possible, tubs were taken out of the fridge up to two minutes before they were entered in the scanner room or presented to participants after scanning. In addition, fresh tubs were presented on each hand within each session.

#### MRI data acquisition

Data were collected using a 3T Siemens Prisma MRI scanner equipped with a 64-channel. High-resolution T1-weighted structural images were acquired for all participants (repetition time (TR) = 2100 ms; echo time (TE) = 2.26 ms voxel size = 1 x 1 x 1 mm). For fMRI, we employed a multi-echo, multi-band echo planar imaging (EPI) acquisition sequence with the following parameters: TR = 1500 ms; TE1= 11 ms, TE2 = 27.25 ms, TE3 = 43.5 ms , TE4 = 59.75 ms; 51 slices with a thickness of 2.60 mm; voxel size = 2.6 x 2.6 x 2.6 mm; field of view 220 mm2, flip angle 77°; 229 volumes; total scanning time = ∼6 minutes. Slices were acquired in interleaved order. To perform fieldmap distorition corrections, a separate field map scan was acquired at the end of each functional scan (TR = 590 ms; TE1= 4.92 ms, TE2 = 7.38 ms; 51 slices with a thickness of 2.60 mm; voxel size = 2.6 x 2.6 x 2.6 mm; field of view 220 mm2, flip angle 46°; interleaved acquisition).

#### MRI preprocessing

Following an initial visual inspection of the data for any abnormalities and artefacts, preprocessing of the fMRI involved several steps. First, motion correction was performed using MCFLIRT [65] from FSL [31] to correct for head movements within the fMRI time series. Fieldmap correction was then applied using the FUGUE command in FSL FEAT to address geometric distortions caused by magnetic field inhomogeneities. Optimal combination of echoes and denoising was achieved using the open-source tool TEDANA [18], which employs principal component analysis (PCA) [76] and independent component analysis (ICA) [71] to separate and remove noise from multi-echo fMRI data. White matter (WM) and cerebrospinal fluid (CSF) signal regression was then applied, with WM and CSF masks obtained from segmentation of T1-weighted scans using the DARTEL tool on SPM12 (https://www.fil.ion.ucl.ac.uk/spm/doc/manual.pdf), which were previously coregistered to each functional scan separately using FSL FLIRT. Resulting denoised data underwent high-pass filtering with a cut-off frequency of 0.005 Hz using to remove low-frequency drifts. Normalisation to Montreal Neurological Institute (MNI) space was achieved in two steps: first, coregistered and segmented grey matter (GM) masks (calculated separately for each run) were normalised to MNI space applying linear and non-linear warping (using FLIRT and FNIRT, respectively). Next, resulting warping parameters were to the functional images. Finally, spatial smoothing was performed with a 5mm FWHM Gaussian kernel to enhance the signal-to-noise ratio.

### Data analysis

#### Self-report data analysis

In order to assess changes in subjective perceived pain across conditions, numerical rate scale (NRS) measures obtained at the end of each scan were compared via means of a one-way repeated measures analysis of variance (ANOVA) with condition (*baseline, tonic pain, recovery)* as within-subjects factor. For post-hoc pairwise comparisons, Sidak Holm correction was applied. In order to test whether the average NRS reported following tonic cold in the scanner was comparable to the average pain intensity perceived on the left hand outside of the scanner, a paired-samples t-test was conducted between average NRS and average VAS (0 = no pain at all; 100 = worst pain imaginable) for each person.

#### Independent Component Analysis (ICA)

In order to explore wholebrain resting state networks arising at group level across rs-fMRI conditions, we conducted ICA analysis. ICA identifies a number of spatially independent patterns of brain activity, previously defined by the researcher, known as components, which correspond to functionally connected brain networks [14; 67]. Each component consists of a time course and a spatial map, where the time course represents the temporal dynamics of the network’s activity, and the spatial map shows the brain regions involved in that network. Unlike seed-based analysis, which requires a priori selection of regions of interest and is limited to exploring connectivity between predefined areas, ICA is a data-driven approach that can detect multiple networks simultaneously without prior assumptions [80]. We conducted ICA within each condition using the MELODIC tool implemented in FSL [5], setting the number of components to 10. Subsequent dual regression analysis was performed to relate the individual subject data back to the group-level components identified in the ICA analysis. Finally, the group level significance of subject-specific spatial maps was assessed using randomise [78] with threshold-free cluster enhancement (TFCE; 5000 permutations, *p*<0.05).

#### Seed-based functional connectivity (FC) analysis

We explored which parts of the brain exhibited BOLD signals that covariates with our predefined seed region, namely the periaqueductal grey (PAG), in order to examine which brain network were associated with PAG activity across conditions. The PAG plays a central role in regulating the body’s response to pain by acting as a relay of activation the descending pain inhibitory pathways [38; 45; 55; 57; 74]. It was therefore hypothesised that changes in FC between this region and the rest of the brain could provide some insights into neural mechanisms underpinning transition between painful and non-painful states. Seed-based FC analyses were performed in FSL FEAT. For each participant, FC maps were generated for each condition (baseline, tonic pain, recovery) using the PAG as the seed region. FC was assessed by correlating the time series from the seed with time series from all other brain voxels. At the group level, we evaluated differences in FC across conditions using a repeated-measures design within a general linear model (GLM), including condition-specific as well as subject-specific explanatory variables (EVs). Our contrast of interest included pairwise comparisons across sessions (Baseline vs Tonic Pain, Tonic Pain vs Recovery and Baseline vs Recovery). Finally, to further explore the relationship between PAG connectivity and pain perception, we computed correlation between the FC maps obtained during the tonic pain condition and participants’ average NRS within a separate GLM. For all group analyses, significance was assessed using Randomise [78] with TFCE correction (5000 permutations, *p*<0.05).

#### Effective connectivity analysis

Finally, we performed spectral dynamic causal modelling (spDCM) to determine whether the patterns of effective connectivity (i.e., the excitatory or inhibitory influence of one brain region on another) within an a priori defined network could account for variability in BOLD signal fluctuations across rs-fMRI conditions. SpDCM utilises a linear random differential equation:

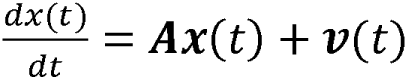

Where **x**(t) denotes the neuronal state vector, **A** represents the matrix of effective connectivity, and **v**(t) captures stochastic endogenous fluctuations. This method focuses on modelling the cross-spectral density G_xy_(ω), derived from the Fourier transform of the cross-covariance function, to capture frequency-domain covariance between neuronal variables. By fitting this model to the observed cross-spectral density, we assessed how effective connectivity explains variations in BOLD fluctuations. For an in-depth explanation of spDCM, see Novelli et al. [52].

Regions of interest (ROIs) were selected according to key loci of the descending pain control pathway, namely the dorsal anterior cingulate cortex (dACC), the anterior insula (AI) the thalamus and the periaqueductal grey (PAG). ROIs were defined using spherical masks based on MNI coordinates obtained from probabilistic maps in the Brainnetome Atlas [21]. Specific map labels, MNI coordinates and radius sizes are detailed in the supplementary information and in Medina and Hughes (2024) [46]. For the periaqueductal gray (PAG), anatomical masks were sourced from the Brainstem Navigator toolkit [7].

SpDCM analyses were conducted in SPM12 [2]. For each participant and fMRI run, a fully connected model was described, where all the possible connections across ROIs were switched on. Model parameters were then estimated via variational Laplace inversion, using the default probability densities provided by SPM12. Parameter estimation (i.e., effective connectivity strengths) was performed at group level within a parametric empirical bayes (PEB) framework [82]. We adopted the same approach described in Hidalgo-Lopez et al., (2021) [26]; first, a PEB design matrix including subject specific regressors in addition to condition specific regressors, to account for repeated measures nature of the design, was constructed. The resulting model evidence decreased compared to the model including condition specific regressors only, and therefore this was the one chosen for final analysis. Thus, to examine differences in effective connectivity within our model across conditions, our final model included 4 regressors: the group mean, the difference between baseline and tonic pain conditions, the difference between tonic pain and recovery conditions and the difference between baseline and recovery conditions. For each regressor, only resulting connections with a posterior probability (the probability of existing) greater than 0.75 are reported (Figure 6).

Finally, we validated our model via leave-one-out cross validation (LOOVC) implemented in SPM12 (spm_dcm_loo.m). For LOOCV, we only included connections where associated parameters showed a pp>0.90 for at least one condition difference, in order to maximise the accuracy in which these connections predicted the conditions.

## Results

### Subjective perceived pain

One participant did not complete the MRI session due to anxiety inside of the scanner. The final sample was N = 29. One-way ANOVA indicated a main effect of Condition (F_(2,56)_= 24.48, *p*<0.001). Pairwise post-hoc comparisons indicated that reported pain was significantly higher following the tonic pain scan (mean(SD) = 17.59(18.23)) than on the other two (baseline mean (SD) = 0.51(1.61); recovery mean (SD) = 2.96(5.15)). Perceived pain following the recovering scan was also significantly greater than in the baseline scan (Figure 2).

**Figure 1.**
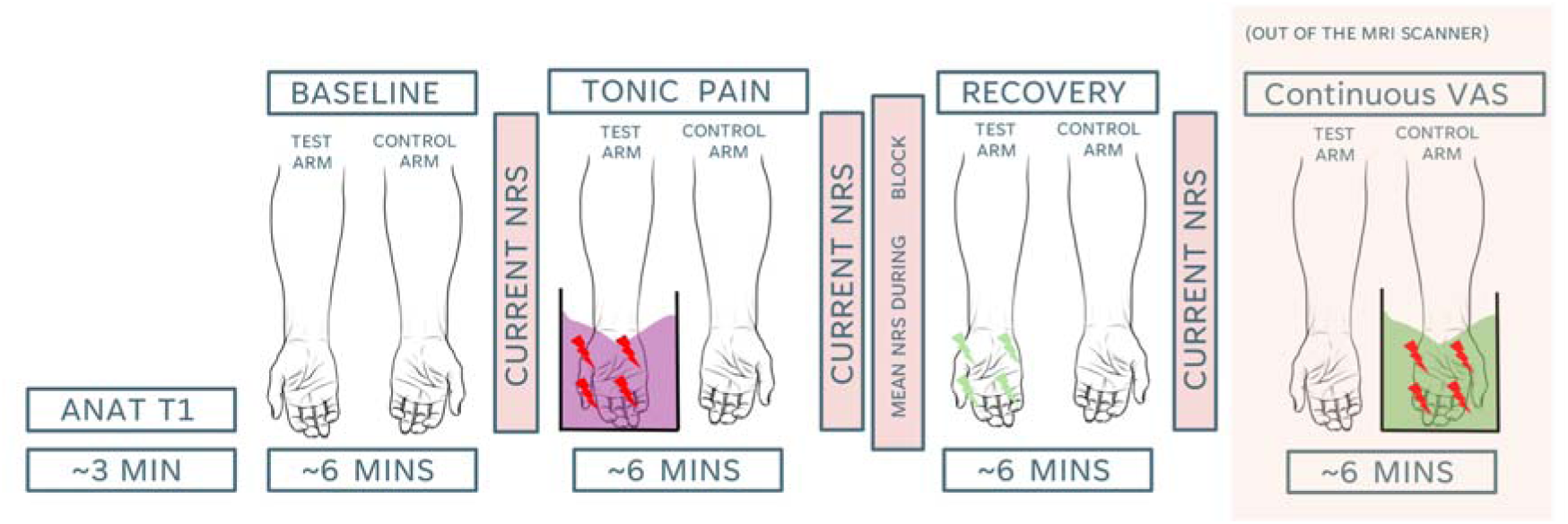
Experimental design. Participants provided their current general perceived levels of pain in a numerical rating scale (NRS) from 0 (‘no pain’) to 100 (‘worst pain imaginable’) following each functional scan. Following the Tonic Pain scan, participants also provided the average perceived pain during tonic cold stimulation in an NRS. During the Tonic Pain condition, all participants immersed their right hand in a gelled-cold water tub. Immediately following fMRI scanning, participants immersed their left hand in a gelled-cold water tub and provided continuous pain intensity ratings in a visual analogue scale (VAS), consisting of a horizontal line anchored from 0 (‘no pain’) to 100 (‘worst pain imaginable’).

**Figure 2.**
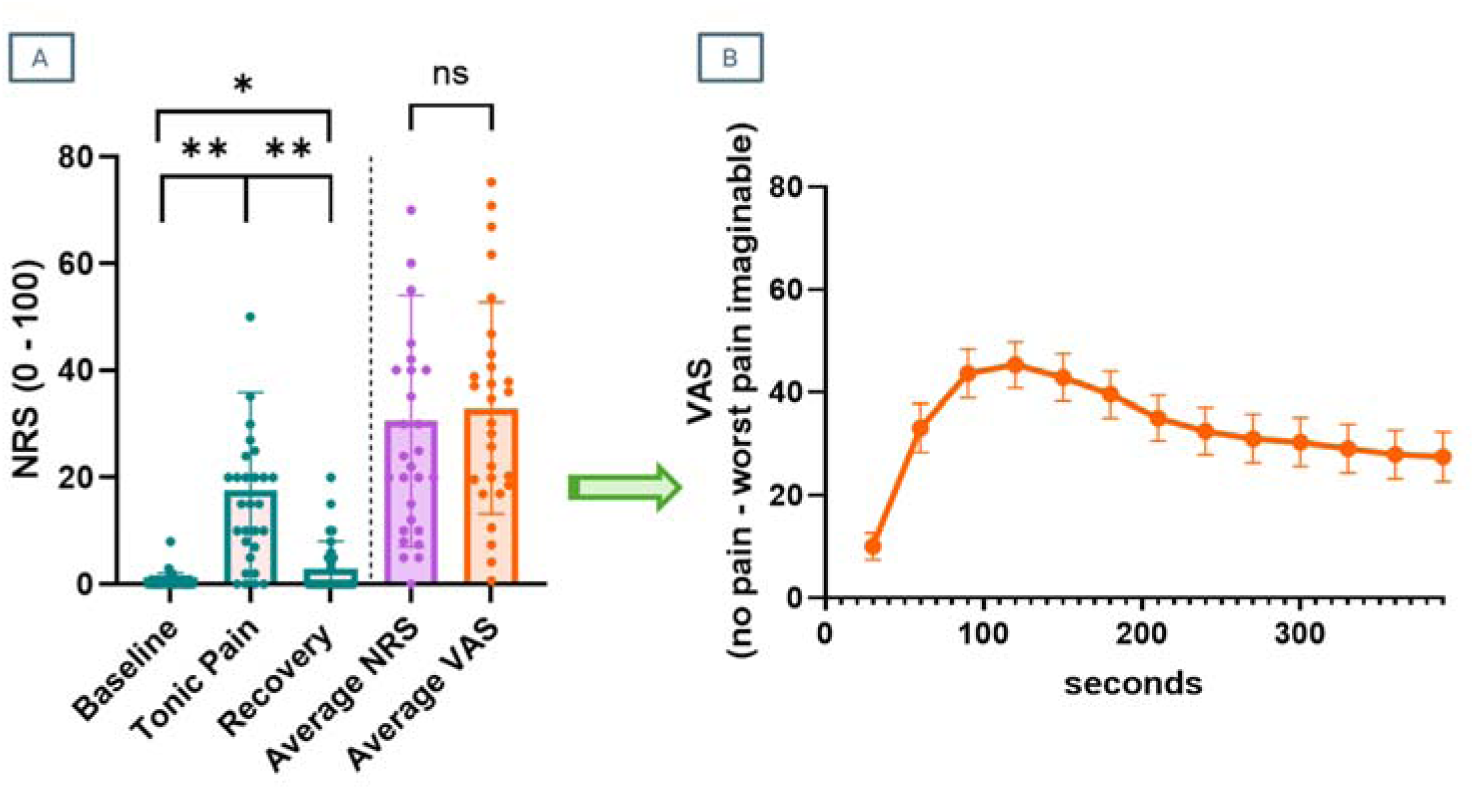
Summary of behavioural results. A) ‘Current pain’ NRS scores were significantly higher following the Tonic Pain condition than on the other two. Average NRS scores about perceived pain intensity estimated by participants during the Tonic Pain scan did not differ from average VAS provided outside of the scanner for tonic cold stimulation on the opposite hand. B) Average continuous VAS ratings for 30-seconds bins across six minutes. Error bars represent standard error of the mean (SEM).

Software for continuous VAS data from one participant failed, and therefore paired-samples t-test included a total N = 28. There was no significant differences between average NRS (mean(SD) = 28.41(20.89)) and computed average VAS (mean(SD) = 32.92(19.75)).

### ICA results

Following ICA, we focused the analysis on components 1-4, which demonstrated distinct spatial patterns relevant to our conditions of interest (Figure 3). Component 1 overlapped considerably across the three conditions and corresponded with the frontal part of the default mode network (DMN) [47; 60]. Component 2 involved sensorimotor regions only during tonic pain and recovery runs, and very reduced areas of frontal and occipital cortices during the baseline run. Component 3 was mostly evident during the tonic pain condition, with activity extending across areas involved in ascending and descending pain control such as dACC, anterior and posterior insulae, basal ganglia, thalamus, midbrain and most of the brainstem. Component 4 involved cerebellar areas overlapping across conditions, extended to lower parts of the brainstem during tonic pain and it included parietal and occipital clusters during baseline and recovery runs. Components 5-10 were considered to be associated with artefactual BOLD activity as well as motion related fluctuations on the edges of the brain and can be found in the Supplementary information b. Following dual regression, TFCE-corrected Z maps indicated that components were statistically significant at group level across conditions (Figure 4).

**Figure 3.**
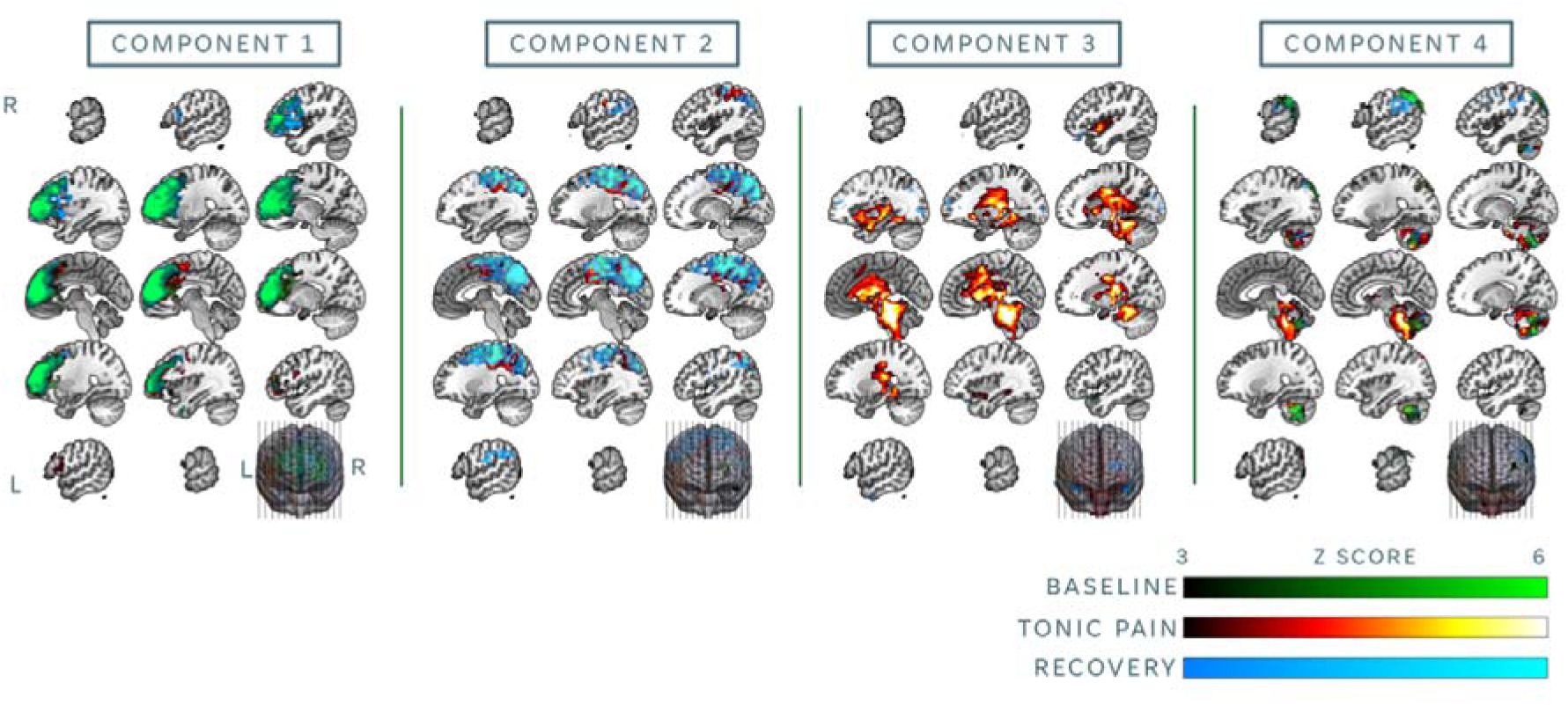
ICA results from components 1-4. Functional networks identified across each condition at group level (green = Baseline, red = Tonic Pain, blue = Recovery). First four components were identified by researchers as the ones capturing variability of interest and not artefactual. L = left. R = right.

**Figure 4.**
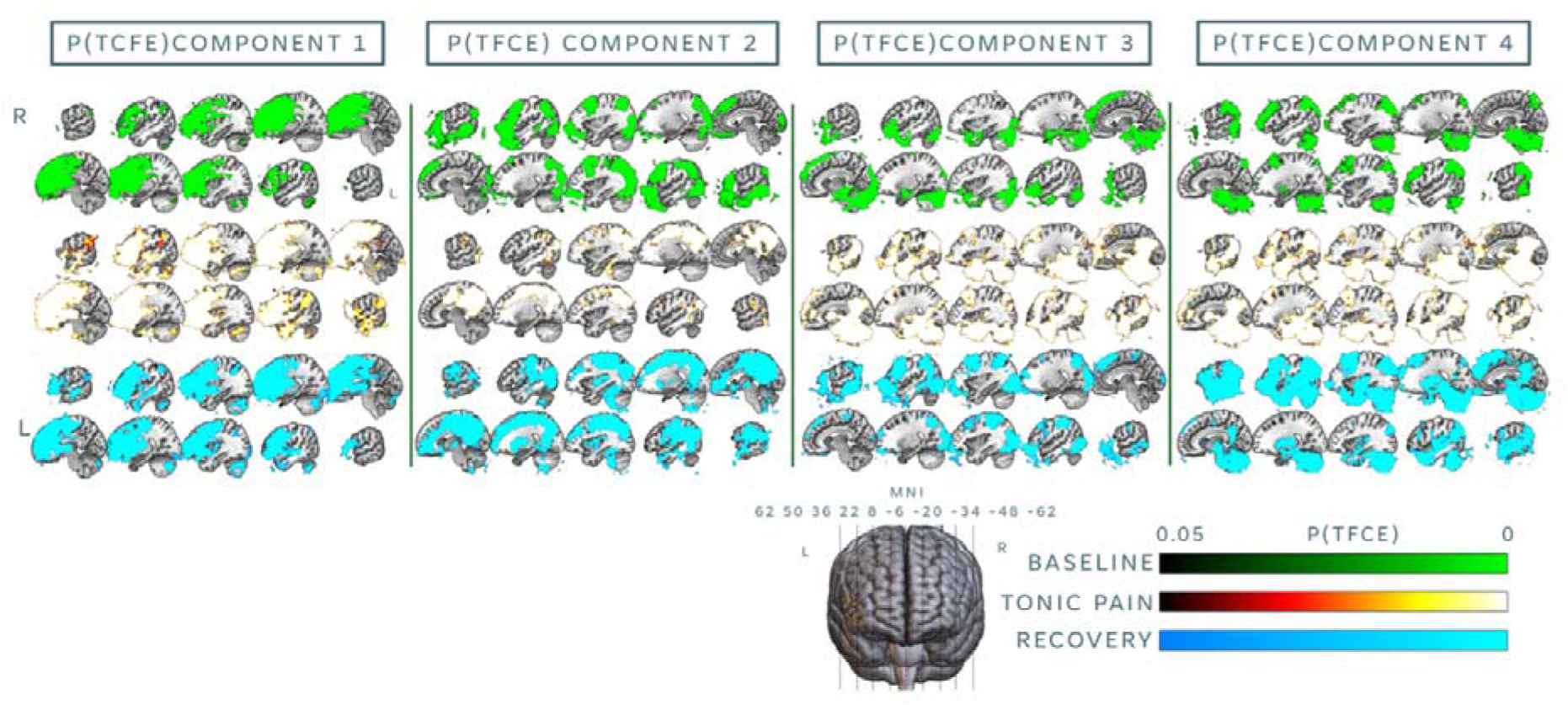
Dual regression results. All maps reflect significant extension of group ICA components at alpha 0.05. TFCE = threshold free cluster enhancement. MNI = Montreal Neurological Institute. L = left. R = right.

### Seed-based FC results

Group analysis revealed that FC between the PAG and widespread cortical and subcortical areas was reduced during tonic pain compared to baseline. Areas involved included the right insula, the intra parietal sulcus (IPS) the ventral ACC, the midcingulate cortex (MCC) and the precuneus and the occipital cortex (Figure 5). We observed no significant FC differences across other contrasts of interest. We also observed no significant correlation between FC of the PAG and the rest of the brain during tonic pain and subjective average NRS scores.

**Figure 5.**
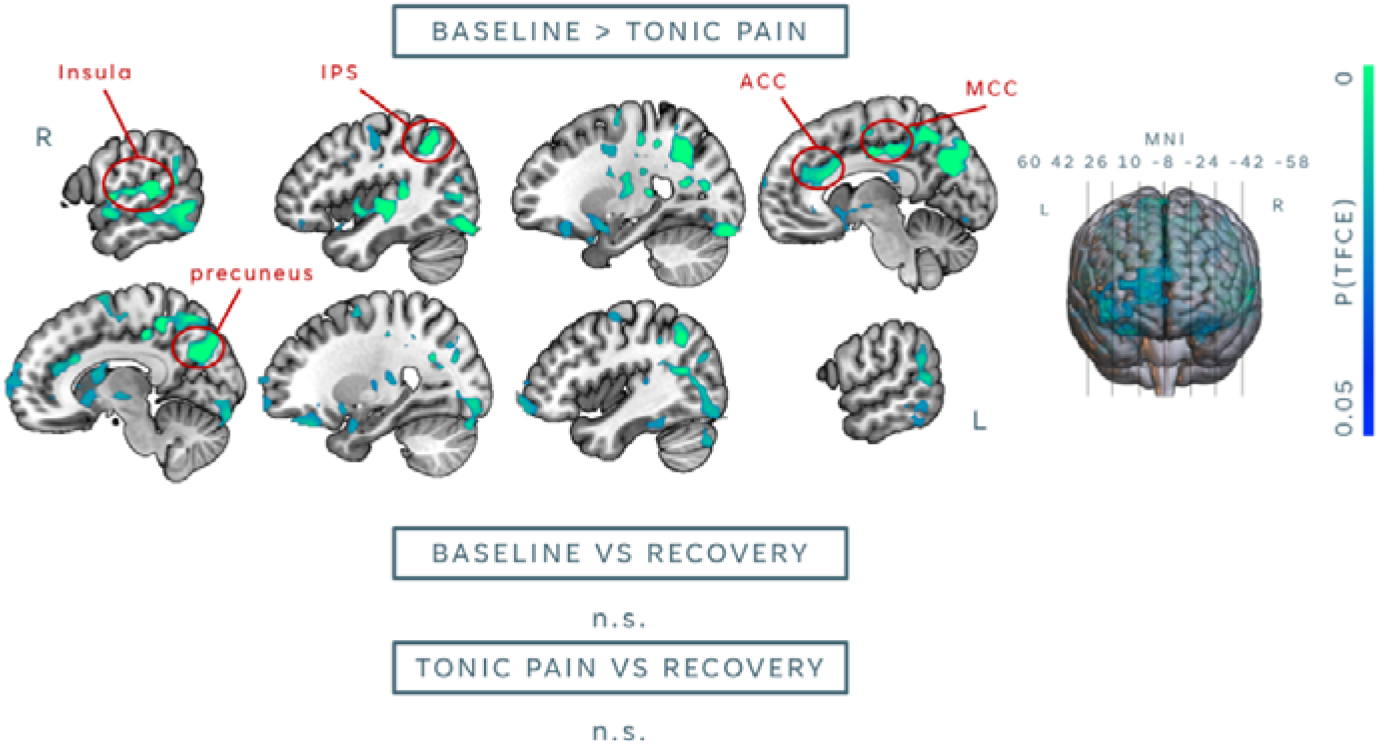
Seed-based connectivity results. Significant map depicts decreases in functional connectivity (FC) with the seed (i.e., PAG) during the Tonic Pain condition compared to Baseline. Results are significant following threshold free cluster enhancement (TFCE) correction at alpha 0.05. IPS = intraparietal sulcus. ACC = anterior cingulate cortex. MCC = midcingulate cortex. MNI = Montreal Neurological Institute. L = left. R = right.

**Figure 6.**
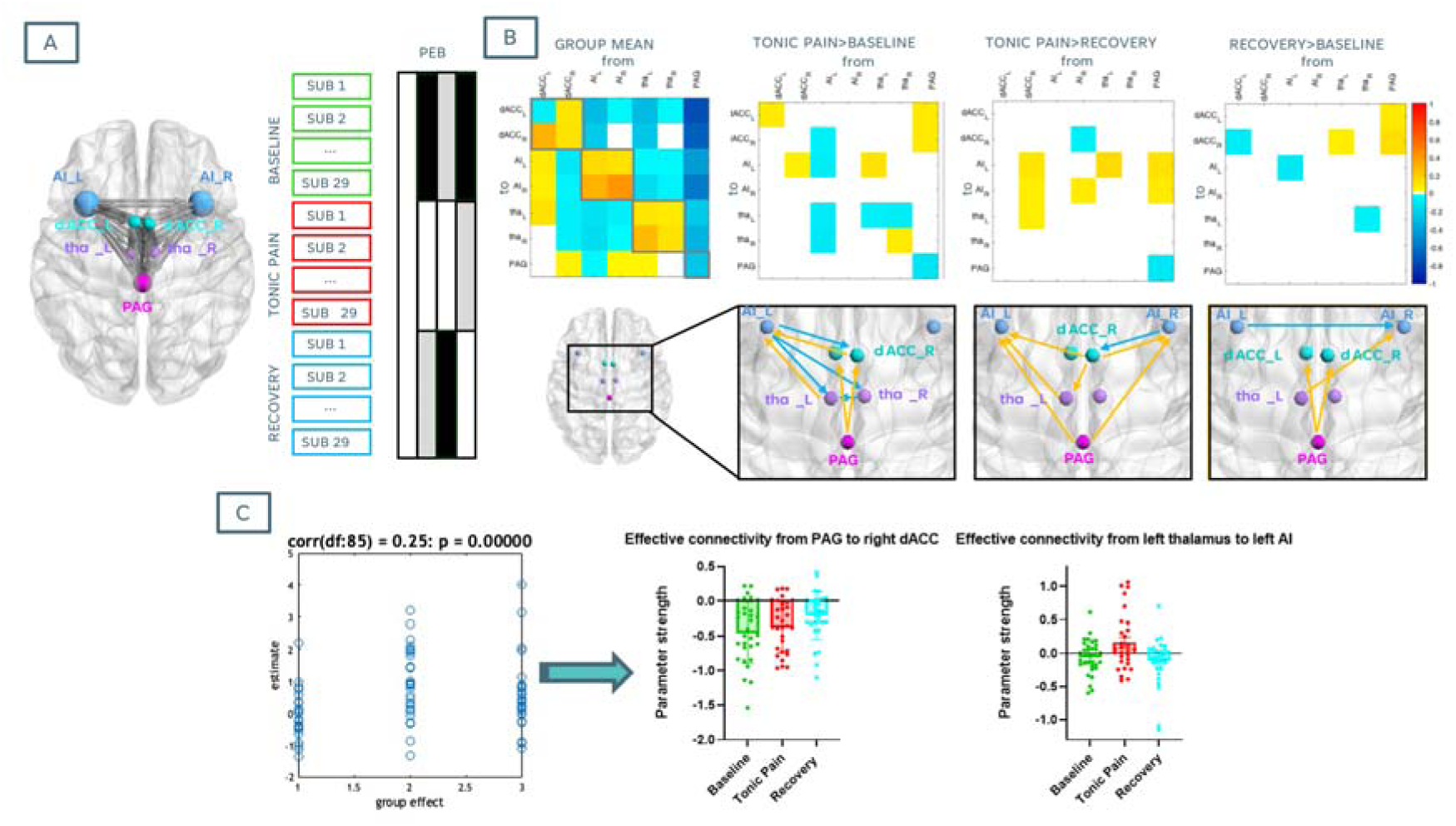
DCM results summary. A) From left to right: Locations of seven nodes comprising the descending pain control pathway used for spDCM analysis. Due to the exploratory nature of the analysis, a fully connected model was initially described. (B) Results from greedy search through nested models; depiction of Parametric Empirical Bayes (PEB) design matrix, comprising scans from all participants across the three experimental conditions (‘Baseline’, ‘Tonic Pain’ and ‘Recovery’). Matrix columns reflect that a group mean, and pairwise comparisons were specified (white colours indicate contrast loads = 1, grey colours indicate contrast loads = 0, black indicates contrast loads = -1). B) Top row shows a depiction of connections with posterior probabilities greater than 0.75 across each contrast of interest. Off-diagonal positive numbers (represented by yellow arrows on the corresponding bottom diagram) represent excitatory connections (*from* locations indicated by rows and *to* locations indicated by columns), and off-diagonal negative numbers (represented by blue arrows on the corresponding bottom diagram) represent inhibitory connections. C) From left to right: Leave –one-out cross-validation (LOOCV) results. Parameters from connections with posterior probabilities greater than 0.9 for pairwise comparison results were used to predict the left-out condition. The correlation between the predicted experimental condition and the actual condition, *r* is the correlation coefficient; connectivity strength across conditions from connections included in LOOCV. AI_L = left anterior insula. AI_R = right anterior insula. dACC_L = left dorsal anterior cingulate cortex. dACC_R = right dorsal anterior cingulate cortex. tha_L = left thalamus. tha_R = right thalamus. PAG = periaqueductal grey.

### DCM results

Individual models per subject and condition achieved acceptable data fit, with percentages of variance explained ranging from 83.88 to 94.89. For PEB results, only parameters for connections showing a posterior probability (i.e., probability of the connection being present vs being absent) greater than 75% are reported and shown in Figure 6.

#### Differences in effective connectivity across conditions

Compared to Baseline, effective connectivity during Tonic Pain was characterised by top-down inhibitory connections from the left AI to right dACC and to thalamus bilaterally, as well as inhibitory activity within the thalamus from right to left ROIs. We also observed bottom-up excitatory connections from the PAG to dACC bilaterally, ads well as from left thalamus and left dACC to the left AI. Within-ROI connectivity strength reflected higher self-modulation of the left dACC and right thalamus and lower self-modulation of the left AI and PAG during tonic pain than at baseline.

Effective connectivity during Tonic Pain compared to Recovery was characterised by bottom-up excitatory activity, specifically from the PAG and from the right dACC to the AI bilaterally, and from the left thalamus to the left AI. We also observed distinct top-down connections, such as inhibitory activity from right AI to right dACC and excitatory activity from the right dACC to the left thalamus. Only the right AI displayed higher self-modulation during Tonic Pain than during Recovery, together with lower self-modulation of the PAG.

Finally, effective connectivity results during the Recovery condition compared to the Baseline condition displayed once again bottom-up excitatory activity from the PAG to the dACC bilaterally, along with excitatory activity from the left thalamus to the right AI. We also observed inhibitory connection from the left AI to the right AI, as well as lower self-modulation of the left AI.

#### Leave-one-out cross-validation (LOOCV)

For validation of our spDCM model, we assessed whether experimental condition could be predicted based on differences in connectivity strength across parameters showing a pp>0.9 in at least one of the contrasts of interest. These connections were from PAG to right dACC and from left thalamus to left AI. LOOCV analysis indicated that these connectivity parameters could successfully predict experimental condition, with Pearson’s correlation between LOO matrix and the predicted condition equal to 0.25 (*p*<0.001) (Figure 6c). Plotting of parameters of interest during LOOCV showed that inhibitory connectivity strength from the PAG to the right dACC progressively decreased across conditions, whereas effective connectivity from the left thalamus to the left AI hovered around zero during Baseline, became on average more excitatory during Tonic Pain, and switched back towards more inhibitory during Recovery.

## Discussion

In this study, we employed a novel strategy to carry out rs-fMRI assessments in healthy individuals before, during and following prolonged tonic noxious cold stimulation, via immersion of the hand in cold gelled-water, mimicking a CPT-like set up. Overall, our experimental paradigm successfully induced varying levels of pain across individuals. Fully exploratory ICA analysis revealed that during Tonic Pain, there was a widespread engagement of areas of the descending pain control network, alongside frontal and sensorimotor engagement that extended across six minutes following stimulation. Results from seed-based connectivity analysis of the PAG with the rest of the brain reflected a decrease in FC with similar areas within this descending pain control network during Tonic Pain compared to Baseline, but not during Recovery. Results from DCM analysis within key nodes of the ascending and descending pain pathways revealed that compared to Baseline, effective connectivity patterns during the Tonic Pain conditions were likely to be characterised by top-down inhibitory influences from the left AI and more excitatory influences bottom-up from PAG and thalamus, the latter ones being also similar during Recovery compared to Baseline. In this section, we discuss the potential mechanisms underlying tonic cold pain according to these results.

At group level, we successfully elicited mild-to-moderate tonic cold pain during six minutes inside of the MRI scanner. Continuous VAS data during analogous stimulation outside of the scanner revealed that subjective pain ratings peaked between 2-3 minutes of noxious stimulation, reducing then to a lower and steady pain sensation. These results are completely consistent with earlier characterisation of responses to the CPT across minutes [11; 16; 19] albeit some variability [68], suggesting that immersion of the hand in gelled-cold water can effectively replicate CPT-like responses. While we did not collect continuous VAS ratings inside of the MR scanner, as this would have compounded the resting state nature of the paradigm, the average NRS ratings provided by participants following the Tonic Pain scan did not significantly differ from average VAS ratings. This result points towards successful reproducibility ratings across both hands, however, future investigations may elucidate the test-retest reliability of tonic cold pain ratings via gelled-cold water.

Crucially, ICA results indicated that, independently to subjective pain, our tonic cold paradigm induced temporally synchronised activity of dACC, midcingulate cortex, AI bilaterally, basal ganglia, thalamus, amygdala, brainstem (including pons, PAG and rostroventral medulla (RVM)) and cerebellum, evidencing engagement of areas across cortical and subcortical descending pain control pathways [3; 4; 12]. This was accompanied by an engagement of widespread sensory and motor areas; while somatosensory activation during stimulation can be usually attributed to processing of discriminative aspects of noxious stimulation, such as location, quality and intensity [54; 73], activation of motor regions during tonic noxious stimulation have been suggested to arise as a result of self-protective aversive reactions during tonic muscle pain [64]. Somatosensory alongside prefrontal engagement was also observed during tonic pain, consistent with previous findings within experimental pain conditions, and suggested to occur to integrate sensory information, allowing to plan and execute appropriate motor responses [15; 50]. Consequently, reappraisal of sensorimotor integration following prolonged tonic cold pain could explain why both prefrontal and sensorimotor independent components persisted during the recovery phase. Another intriguing finding from our ICA analysis was the increased extension of the brainstem-cerebellar component during tonic noxious stimulation compared to the other conditions. This is consistent with positron emission tomography (PET) findings during the CPT, where brainstem activation was related to initial responses to cold noxious stimulation and changes in galvanic skin responses, suggesting its involvement in sympathetic responses during noxious cold [59]. Increases in regional cerebral blood flow (rCBF) have also been observed in across descending pain control pathway and cerebellum during the CPT [10]. Increasing evidence suggests that the cerebellum plays an crucial modulatory role during nociception [49], temperature detection [84] and top-down pain modulation via its communication with sensorimotor, executive, reward and limbic regions through the brainstem [40]. Taken together, these findings indicate that our cold pain paradigm successfully elicited functional connectivity patterns consistent with those observed in studies of perceived pain generally, and the CPT specifically. Notably, these results emerged from an entirely data-driven approach, such as ICA, which minimises bias in the identification of neural networks. This underscores the utility of the current paradigm as a robust method for inducing CPT-like pain neural signatures during fMRI in healthy individuals, offering a viable platform for investigating pain processing and connectivity dynamics.

Our seed-based connectivity analysis revealed a disengagement of the PAG with an array of cortical areas, including right insula, IPS, ACC, midcingulate cortex, precuneus and occipital cortex, during tonic pain compared to baseline, partly overlapping with previous findings in healthy volunteers [43]. This was likely due to a shift towards an engagement with downstream interactions with the brainstem and spinal cord to inhibit cold noxious input [39; 44; 79]. While previous findings showed correlations between PAG activation during cold tonic [36] as well as evoked [48] pain and subjective ratings, FC of the PAG did not correlate with individual subjective pain scores; this result suggests that PAG-led pain modulation might rely more on the action of top-down PAG-RVM-spinal opioidergic pathways than on the extent of PAG-brain synchronisation, which has been suggested to be more closely related to autonomic regulation during pain [27; 44]. Nevertheless, sympathetic regulation may indirectly modulate PAG-RVM inhibition through stress induced analgesia [38]. Moreover, PAG-cortex disengagement was not observed during the Recovery condition compared to the Tonic Pain condition, hinting towards sustained modulatory action following withdrawal of tonic cold stimuli.

In fact, our spDCM results revealed that effective connectivity patterns during the Tonic Pain condition, compared to Baseline, were most likely characterised by more excitatory influences from the PAG to the dACC bilaterally and from the thalamus to the AI, a pattern that persisted during Recovery compared to Baseline. The PAG is a central hub for pain modulation, acting top-down and as a bottom-up relay of pain responses and nociceptive drive, respectively (for a review, see Linnman et al. [41]). During tonic pain, it is reasonable to argue that the PAG might amplify excitatory input to the dACC to enhance the perception and emotional salience and aversion of pain, encouraging protective or adaptive behaviours [83]. This would also explain why the Tonic Pain condition was characterised by heightened excitatory influences from the PAG to the AI than during Recovery. Nevertheless, bottom-up PAG-dACC excitatory activity persisted during Recovery compared to Baseline, which suggests recalibration of both sensory and affective circuits [27; 83]. The PAG might sustain excitatory inputs to the dACC during early recovery to process and resolve the lingering affective and evaluative aspects of pain. Similarly, our results suggest that the left thalamus acted as a relay of nociceptive drive to the left AI during tonic pain, probably resulting from the activation of spinothalamic-cortical pathways [6; 22; 23; 29]. Our LOOCV results showed that this influenced switched towards inhibition during the Recovery scan, arguably reflecting and adaptative action to recalibrate the pain network and reset to baseline. Conversely, compared to Baseline, effective connectivity during tonic pain was likely characterised by top-down inhibitory action of the AI on the dACC and thalamus bilaterally, which might reflect a regulatory attempt to downscale the dACC’s activity and a top-down mechanism to prevent excessive engagement with the aversive aspects of pain [1; 56], which could otherwise lead to heightened distress or maladaptive behavioural responses [8].

In general, LOOCV revealed that effective connectivity of the PAG-dACC and thalamus-AI pathways could successfully predict experimental conditions, highlighting the state-specific modulation of pain-related networks. These findings underscore the mechanistic roles of ascending pathways in processing the sensory, affective, and regulatory components of pain and its resolution. The predictive accuracy of these connections positions them as potential biomarkers for characterising the dynamics of pain states and transitions, with implications for personalised pain management strategies.

## Conclusion

In conclusion, this study establishes the cold gelled-water paradigm as a viable in-scanner alternative to the CPT, effectively replicating its neurophysiological signatures and enabling the study of tonic pain in healthy individuals. We revealed distinct neural dynamics underpinning CPT-like pain, including PAG-driven excitatory inputs to dACC and AI that amplified pain salience, alongside AI-mediated inhibitory control over dACC and thalamus to regulate sustained nociceptive and affective responses. Each analysis method provided unique insights: ICA captured large-scale network dynamics, seed-based connectivity revealed specific functional reorganisations of the PAG, and DCM clarified directional influences within pain pathways. Together, these findings underscore the paradigm’s utility for probing pain modulation and transitions, offering a robust platform for studying adaptive and maladaptive neural responses to pain.

## Acknowledgements

This study was funded through an Academy of Medical Sciences Springboard grant (SBF007\100108). We would like to thank Jamie Roberts for his assistance with the scanning protocol. We would like to express our sincere gratitude to all of our study participants. The authors have no conflicts of interest.

## Data Availability

The data that support the findings of this study are available from the corresponding author upon reasonable request. Scripts for data preprocessing and analysis are available on osf.io/txrwm/files/osfstorage.

## References

[1] Alonso-Matielo H, Zhang Z, Gambeta E, Huang J, Chen L, de Melo GO, Dale CS, Zamponi GW. Inhibitory insula-ACC projections modulate affective but not sensory aspects of neuropathic pain. Molecular Brain 2023;16(1):64.

[2] Ashburner J, Barnes G, Chen C-C, Daunizeau J, Flandin G, Friston K, Kiebel S, Kilner J, Litvak V, Moran R. SPM12 manual. Wellcome Trust Centre for Neuroimaging, London, UK 2014;2464(4).

[3] Bannister K. Descending pain modulation: influence and impact. Current Opinion in Physiology 2019;11:62–66.

[4] Bannister K, Dickenson A. The plasticity of descending controls in pain: translational probing. The Journal of physiology 2017;595(13):4159–4166.

[5] Beckmann CF, Smith SM. Probabilistic independent component analysis for functional magnetic resonance imaging. IEEE transactions on medical imaging 2004;23(2):137–152.

[6] Benarroch EE, Benarroch EE. 674Central Processing and Modulation of Pain. In: EE Benarroch, editor. Neuroscience for Clinicians: Basic Processes, Circuits, Disease Mechanisms, and Therapeutic Implications: Oxford University Press, 2021. p. 0.

[7] Bianciardi M. Brainstem Navigator [Tool/Resource.]. Washington: NITRC, 2021.

[8] Buchmann J, Baumann N, Meng K, Semrau J, Kuhl J, Pfeifer K, Kazén M, Vogel H, Faller H. Endurance and avoidance response patterns in pain patients: Application of action control theory in pain research. PLoS One 2021;16(3):e0248875.

[9] Carlsson AM. Assessment of chronic pain. I. Aspects of the reliability and validity of the visual analogue scale. Pain 1983;16(1):87–101.

[10] Casey KL, Minoshima S, Morrow TJ, Koeppe RA. Comparison of human cerebral activation pattern during cutaneous warmth, heat pain, and deep cold pain. Journal of neurophysiology 1996;76(1):571–581.

[11] Chang PF, Arendt-Nielsen L, Chen AC. Dynamic changes and spatial correlation of EEG activities during cold pressor test in man. Brain research bulletin 2002;57(5):667–675.

[12] Chen Q, Heinricher MM. Descending Control Mechanisms and Chronic Pain. Current Rheumatology Reports 2019;21(5):13.

[13] Clewett D, Schoeke A, Mather M. Amygdala functional connectivity is reduced after the cold pressor task. Cognitive, Affective, & Behavioral Neuroscience 2013;13:501–518.

[14] Damoiseaux JS, Rombouts SA, Barkhof F, Scheltens P, Stam CJ, Smith SM, Beckmann CF. Consistent resting-state networks across healthy subjects. Proceedings of the national academy of sciences 2006;103(37):13848–13853.

[15] Dancey E, Murphy BA, Andrew D, Yielder P. The effect of local vs remote experimental pain on motor learning and sensorimotor integration using a complex typing task. PAIN 2016;157(8):1682–1695.

[16] Devoize L, Chalaye P, Lafrenaye S, Marchand S, Dallel R. Relationship between adaptation and cardiovascular response to tonic cold and heat pain adaptability to tonic pain and cardiovascular responses. European Journal of Pain 2016;20(5):731–741.

[17] Duncko R, Johnson L, Merikangas K, Grillon C. Working memory performance after acute exposure to the cold pressor stress in healthy volunteers. Neurobiology of learning and memory 2009;91(4):377–381.

[18] DuPre E, Salo T, Ahmed Z, Bandettini PA, Bottenhorn KL, Caballero-Gaudes C, Dowdle LT, Gonzalez-Castillo J, Heunis S, Kundu P. TE-dependent analysis of multi-echo fMRI with* tedana. Journal of Open Source Software 2021;6(66):3669.

[19] Eccleston C. The attentional control of pain: methodological and theoretical concerns. Pain 1995;63(1):3–10.

[20] Elias SO, Ajayi RE. Effect of sympathetic autonomic stress from the cold pressor test on left ventricular function in young healthy adults. Physiological reports 2019;7(2):e13985.

[21] Fan L, Li H, Zhuo J, Zhang Y, Wang J, Chen L, Yang Z, Chu C, Xie S, Laird AR. The human brainnetome atlas: a new brain atlas based on connectional architecture. Cerebral cortex 2016;26(8):3508–3526.

[22] Gélébart J, Garcia-Larrea L, Frot M. Amygdala and anterior insula control the passage from nociception to pain. Cerebral Cortex 2022;33(7):3538–3547.

[23] Groh A, Mease R, Krieger P. Pain processing in the thalamocortical system. e-Neuroforum 2017;23(3):117–122.

[24] Grouper H, Löffler M, Flor H, Eisenberg E, Pud D. Increased functional connectivity between limbic brain areas in healthy individuals with high versus low sensitivity to cold pain: A resting state fMRI study. Plos one 2022;17(4):e0267170.

[25] Hendriks-Balk MC, Megdiche F, Pezzi L, Reynaud O, Da Costa S, Bueti D, Van De Ville D, Wuerzner G. Brainstem correlates of a cold pressor test measured by ultra-high field fMRI. Frontiers in neuroscience 2020;14:39.

[26] Hidalgo-Lopez E, Zeidman P, Harris T, Razi A, Pletzer B. Spectral dynamic causal modelling in healthy women reveals brain connectivity changes along the menstrual cycle. Communications Biology 2021;4(1):954.

[27] Hohenschurz-Schmidt D, Calcagnini G, Dipasquale O, Jackson J, Medina S, O’Daly O, O’Muircheartaigh J, de Lara R, Williams S, McMahon S. Linking pain sensation to the autonomic nervous system: the role of the anterior cingulate and periaqueductal gray resting-state networks. Front Neurosci. 2020. Importance of the autonomic nervous system in pain states 2020.

[28] Huang M, Yoo J-K, Stickford AS, Moore JP, Hendrix JM, Crandall CG, Fu Q. Early sympathetic neural responses during a cold pressor test linked to pain perception. Clinical Autonomic Research 2021;31:215–224.

[29] Jalon I, Berger A, Shofty B, Goldway N, Artzi M, Gurevitch G, Hochberg U, Tellem R, Hendler T, Gonen T. Lesions to both somatic and affective pain pathways lead to decreased salience network connectivity. Brain 2023;146(5):2153–2162.

[30] Jarrahi B, Martucci KT, Nilakantan AS, Mackey S. Cold water pressor test differentially modulates functional network connectivity in fibromyalgia patients compared with healthy controls, Proceedings of the 2018 40th Annual International Conference of the IEEE Engineering in Medicine and Biology Society (EMBC): IEEE, 2018. pp. 578–582.

[31] Jenkinson M, Beckmann CF, Behrens TE, Woolrich MW, Smith SM. Fsl. Neuroimage 2012;62(2):782–790.

[32] Johnson MH, Petrie SM. The effects of distraction on exercise and cold pressor tolerance for chronic low back pain sufferers. Pain 1997;69(1-2):43–48.

[33] Kasch H, Qerama E, Bach FW, Jensen TS. Reduced cold pressor pain tolerance in non-recovered whiplash patients: a 1-year prospective study. European Journal of Pain 2005;9(5):561–569.

[34] Kennedy DL, Kemp HI, Ridout D, Yarnitsky D, Rice AS. Reliability of conditioned pain modulation: a systematic review. Pain 2016;157(11):2410–2419.

[35] King M, Carnahan H. Revisiting the brain activity associated with innocuous and noxious cold exposure. Neuroscience & Biobehavioral Reviews 2019;104:197–208.

[36] La Cesa S, Tinelli E, Toschi N, Di Stefano G, Collorone S, Aceti A, Francia A, Cruccu G, Truini A, Caramia F. fMRI pain activation in the periaqueductal gray in healthy volunteers during the cold pressor test. Magnetic resonance imaging 2014;32(3):236–240.

[37] Lapotka M, Ruz M, Salamanca Ballesteros A, Ocón Hernández O. Cold pressor gel test: A safe alternative to the cold pressor test in fMRI. Magnetic Resonance in Medicine 2017;78(4):1464–1468.

[38] Lau BK, Vaughan CW. Descending modulation of pain: the GABA disinhibition hypothesis of analgesia. Current opinion in neurobiology 2014;29:159–164.

[39] Leith JL, Koutsikou S, Lumb BM, Apps R. Spinal processing of noxious and innocuous cold information: differential modulation by the periaqueductal gray. Journal of Neuroscience 2010;30(14):4933–4942.

[40] Li CN, Keay KA, Henderson LA, Mychasiuk R. Re-examining the Mysterious Role of the Cerebellum in Pain. Journal of Neuroscience 2024;44(17).

[41] Linnman C, Moulton EA, Barmettler G, Becerra L, Borsook D. Neuroimaging of the periaqueductal gray: state of the field. Neuroimage 2012;60(1):505–522.

[42] Luo J, Zhu H-Q, Gou B, Wang X-Q. Neuroimaging assessment of pain. Neurotherapeutics 2022;19(5):1467–1488.

[43] Makovac E, Dipasquale O, Jackson JB, Medina S, O’Daly O, O’Muircheartaigh J, de Lara Rubio A, Williams SC, McMahon SB, Howard MA. Sustained perturbation in functional connectivity induced by cold pain. European Journal of Pain 2020;24(9):1850–1861.

[44] Makovac E, Venezia A, Hohenschurz-Schmidt D, Dipasquale O, Jackson JB, Medina S, O’Daly O, Williams SC, McMahon SB, Howard MA. The association between pain-induced autonomic reactivity and descending pain control is mediated by the periaqueductal grey. The Journal of Physiology 2021;599(23):5243–5260.

[45] Mason P. Central mechanisms of pain modulation. Current opinion in neurobiology 1999;9(4):436–441.

[46] Medina S, Hughes S. Immersion in nature attenuates the development of mechanical secondary hyperalgesia: a role for insulo-thalamic effective connectivity. bioRxiv 2024:2024.2010. 2011.617804.

[47] Menon V. 20 years of the default mode network: A review and synthesis. Neuron 2023;111(16):2469–2487.

[48] Mohr C, Leyendecker S, Mangels I, Machner B, Sander T, Helmchen C. Central representation of cold-evoked pain relief in capsaicin induced pain: an event-related fMRI study. Pain 2008;139(2):416–430.

[49] Moulton EA, Schmahmann JD, Becerra L, Borsook D. The cerebellum and pain: passive integrator or active participator? Brain research reviews 2010;65(1):14–27.

[50] Nickel MM, Ta Dinh S, May ES, Tiemann L, Hohn VD, Gross J, Ploner M. Neural oscillations and connectivity characterizing the state of tonic experimental pain in humans. Human Brain Mapping 2020;41(1):17–29.

[51] Nouwen A, Cloutier C, Kappas A, Warbrick T, Sheffield D. Effects of focusing and distraction on cold pressor–induced pain in chronic back pain patients and control subjects. The Journal of Pain 2006;7(1):62–71.

[52] Novelli L, Friston K, Razi A. Spectral dynamic causal modeling: A didactic introduction and its relationship with functional connectivity. Network Neuroscience 2024;8(1):178–202.

[53] Oaks Z, Stage A, Middleton B, Faraone SV, Johnson B. Clinical utility of the cold pressor test: evaluation of pain patients and treatment of opioid-induced hyperalgesia and fibromyalgia with low dose naltrexone. Discovery Medicine 2018;26(144):197–206.

[54] Oshiro Y, Quevedo AS, McHaffie JG, Kraft RA, Coghill RC. Brain mechanisms supporting discrimination of sensory features of pain: a new model. Journal of Neuroscience 2009;29(47):14924–14931.

[55] Ossipov MH. The perception and endogenous modulation of pain. Scientifica 2012;2012(1):561761.

[56] Ossipov MH, Dussor GO, Porreca F. Central modulation of pain. The Journal of clinical investigation 2010;120(11):3779–3787.

[57] Ossipov MH, Morimura K, Porreca F. Descending pain modulation and chronification of pain. Current opinion in supportive and palliative care 2014;8(2):143–151.

[58] Paccione CE, Bruehl S, Diep LM, Rosseland LA, Stubhaug A, Jacobsen HB. The indirect impact of heart rate variability on cold pressor pain tolerance and intensity through psychological distress in individuals with chronic pain: the Tromsø Study. Pain reports 2022;7(2):e970.

[59] Petrovic P, Petersson KM, Hansson P, Ingvar M. Brainstem involvement in the initial response to pain. Neuroimage 2004;22(2):995–1005.

[60] Raichle ME. The brain’s default mode network. Annual review of neuroscience 2015;38(1):433–447.

[61] Richardson HL, Macey PM, Kumar R, Valladares EM, Woo MA, Harper RM. Neural and physiological responses to a cold pressor challenge in healthy adolescents. Journal of neuroscience research 2013;91(12):1618–1627.

[62] Ríos-León M, Demertzis E, Palazón-García R, Taylor J. Tonic Cold Pain Temporal Summation and Translesional Cold Pressor Test-Induced Pronociception in Spinal Cord Injury: Association with Spontaneous and Below-Level Neuropathic Pain, Proceedings of the Healthcare, Vol. 12: MDPI, 2024. p. 2300.

[63] Roatta S, Micieli G, Bosone D, Losano G, Bini R, Cavallini A, Passatore M. Effect of generalised sympathetic activation by cold pressor test on cerebral haemodynamics in healthy humans. Journal of the autonomic nervous system 1998;71(2-3):159–166.

[64] Rossi S, della Volpe R, Ginanneschi F, Ulivelli M, Bartalini S, Spidalieri R, Rossi A. Early somatosensory processing during tonic muscle pain in humans: relation to loss of proprioception and motor ‘defensive’ strategies. Clinical Neurophysiology 2003;114(7):1351–1358.

[65] Saccà V, Saricà A, Novellino F, Barone S, Tallarico T, Filippelli E, Granata A, Valentino P, Quattrone A. Evaluation of the MCFLIRT Correction Algorithm in Head Motion from Resting State fMRI Data. International Journal of Biomedical and Biological Engineering 2018;12(3):58–61.

[66] Safety: ACoM, Greenberg TD, Hoff MN, Gilk TB, Jackson EF, Kanal E, McKinney AM, Och JG, Pedrosa I, Rampulla TL. ACR guidance document on MR safe practices: Updates and critical information 2019. Journal of Magnetic Resonance Imaging 2020;51(2):331–338.

[67] Schöpf V, Kasess C, Lanzenberger R, Fischmeister F, Windischberger C, Moser E. Fully exploratory network ICA (FENICA) on resting-state fMRI data. Journal of neuroscience methods 2010;192(2):207–213.

[68] Shao S, Shen K, Yu K, Wilder-Smith EP, Li X. Frequency-domain EEG source analysis for acute tonic cold pain perception. Clinical Neurophysiology 2012;123(10):2042–2049.

[69] Smitha K, Akhil Raja K, Arun K, Rajesh P, Thomas B, Kapilamoorthy T, Kesavadas C. Resting state fMRI: A review on methods in resting state connectivity analysis and resting state networks. The neuroradiology journal 2017;30(4):305–317.

[70] Spielberger CD, Gonzalez-Reigosa F, Martinez-Urrutia A, Natalicio LF, Natalicio DS. The state-trait anxiety inventory. Revista Interamericana de Psicologia/Interamerican journal of psychology 1971;5(3 & 4).

[71] Steel A, Garcia BD, Silson EH, Robertson CE. Evaluating the efficacy of multi-echo ICA denoising on model-based fMRI. Neuroimage 2022;264:119723.

[72] Stevens A, Batra A, Kötter I, Bartels M, Schwarz J. Both pain and EEG response to cold pressor stimulation occurs faster in fibromyalgia patients than in control subjects. Psychiatry research 2000;97(2-3):237–247.

[73] Sun G, McCartin M, Liu W, Zhang Q, Kenefati G, Chen ZS, Wang J. Temporal pain processing in the primary somatosensory cortex and anterior cingulate cortex. Molecular Brain 2023;16(1):3.

[74] Tobaldini G, Sardi NF, Guilhen VA, Fischer L. Pain inhibits pain: an ascending-descending pain modulation pathway linking mesolimbic and classical descending mechanisms. Molecular neurobiology 2019;56:1000–1013.

[75] Vaegter HB, Handberg G, Graven-Nielsen T. Hypoalgesia after exercise and the cold pressor test is reduced in chronic musculoskeletal pain patients with high pain sensitivity. The Clinical journal of pain 2016;32(1):58–69.

[76] Viviani R, Grön G, Spitzer M. Functional principal component analysis of fMRI data. Human brain mapping 2005;24(2):109–129.

[77] Watso JC, Huang M, Belval LN, Cimino 3rd FA, Jarrard CP, Hendrix JM, Hinojosa-Laborde C, Crandall CG. Low-dose fentanyl reduces pain perception, muscle sympathetic nerve activity responses, and blood pressure responses during the cold pressor test. American Journal of Physiology-Regulatory, Integrative and Comparative Physiology 2022;322(1):R64–R76.

[78] Winkler AM, Ridgway GR, Webster MA, Smith SM, Nichols TE. Permutation inference for the general linear model. NeuroImage 2014;92:381–397.

[79] Xin L, Geller EB, Liu-Chen L-Y, Chen C, Adler MW. Substance P release in the rat periaqueductal gray and preoptic anterior hypothalamus after noxious cold stimulation: effect of selective mu and kappa opioid agonists. Journal of Pharmacology and Experimental Therapeutics 1997;282(2):1055–1063.

[80] Yang J, Gohel S, Vachha B. Current methods and new directions in resting state fMRI. Clinical Imaging 2020;65:47–53.

[81] Zacny JP, Coalson DW, Young CJ, Klafta JM, Lichtor JL, Rupani G, Thapar P, Apfelbaum JL. Propofol at conscious sedation doses produces mild analgesia to cold pressor-induced pain in healthy volunteers. Journal of clinical anesthesia 1996;8(6):469–474.

[82] Zeidman P, Jafarian A, Seghier ML, Litvak V, Cagnan H, Price CJ, Friston KJ. A guide to group effective connectivity analysis, part 2: Second level analysis with PEB. Neuroimage 2019;200:12–25.

[83] Zhang H, Zhu Z, Ma W-X, Kong L-X, Yuan P-C, Bu L-F, Han J, Huang Z-L, Wang Y-Q. The contribution of periaqueductal gray in the regulation of physiological and pathological behaviors. Frontiers in Neuroscience 2024;18:1380171.

[84] Zunhammer M, Busch V, Griesbach F, Landgrebe M, Hajak G, Langguth B. rTMS over the cerebellum modulates temperature detection and pain thresholds through peripheral mechanisms. Brain stimulation 2011;4(4):210–217. e211.

