## Supplementary Information for "Neural Dynamics of Tonic Cold Pain: A Novel Investigation of an In-Scanner Alternative to the Cold Pressor Test in Healthy Individuals"

a. **ROI definition**

| **ROI** | **BNA atlas labels** | **MNI coordinates** | | **Radius size** |
| --- | --- | --- | --- | --- |
|  |  | **Left** | **Right** |  |
| dorsal ACC | A24cd, caudodorsal area 24 | -5, 7, 37 | 4, 6, 38 | 4mm |
| AI | dIa, dorsal agranular insula | -34, 18, 1 | 36, 18, 1 | 8mm |
| thalamus | mPFtha, medial pre-frontal thalamus | -7, -12, 5 | 7, -11, 6 | 4mm |

**Supplementary table 1.** ROI definition details. ACC = anterior cingulate cortex; AI = anterior insula; BNA = Brainnetome Atlas; MNI = Montreal Neurological Institute

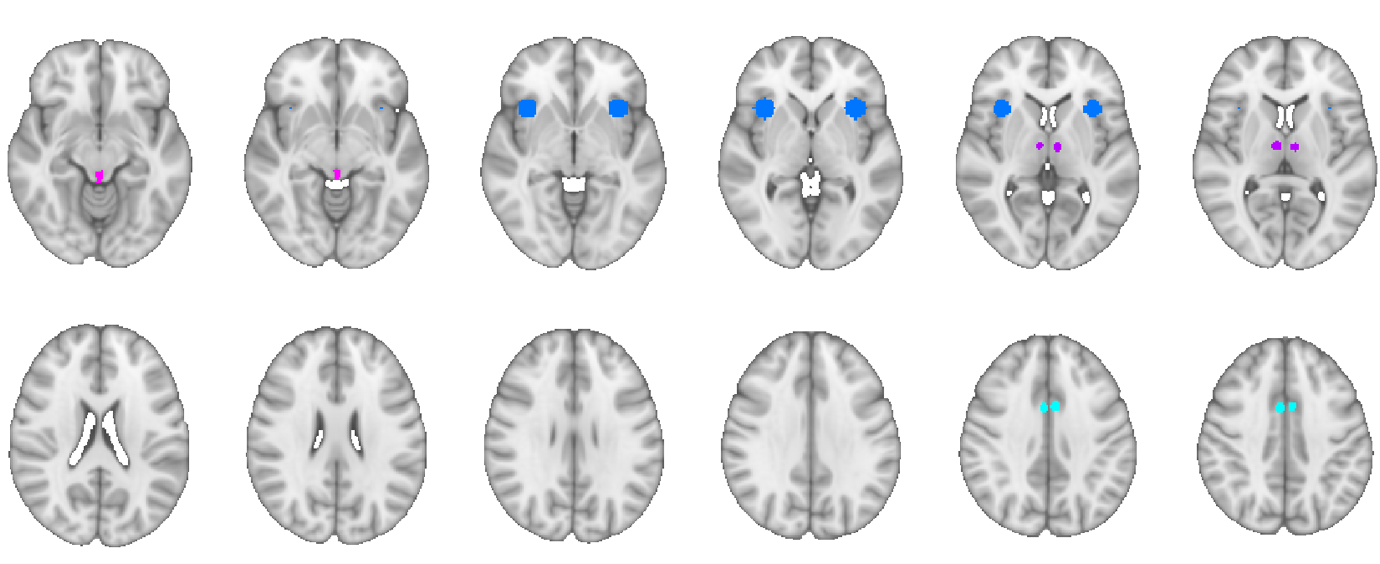

**Supplementary Figure 1.** Depiction of dACC (light blue), AI (dark blue), thalamus (purple) and PAG (pink) ROIs.

b. **ICA components 5-10**

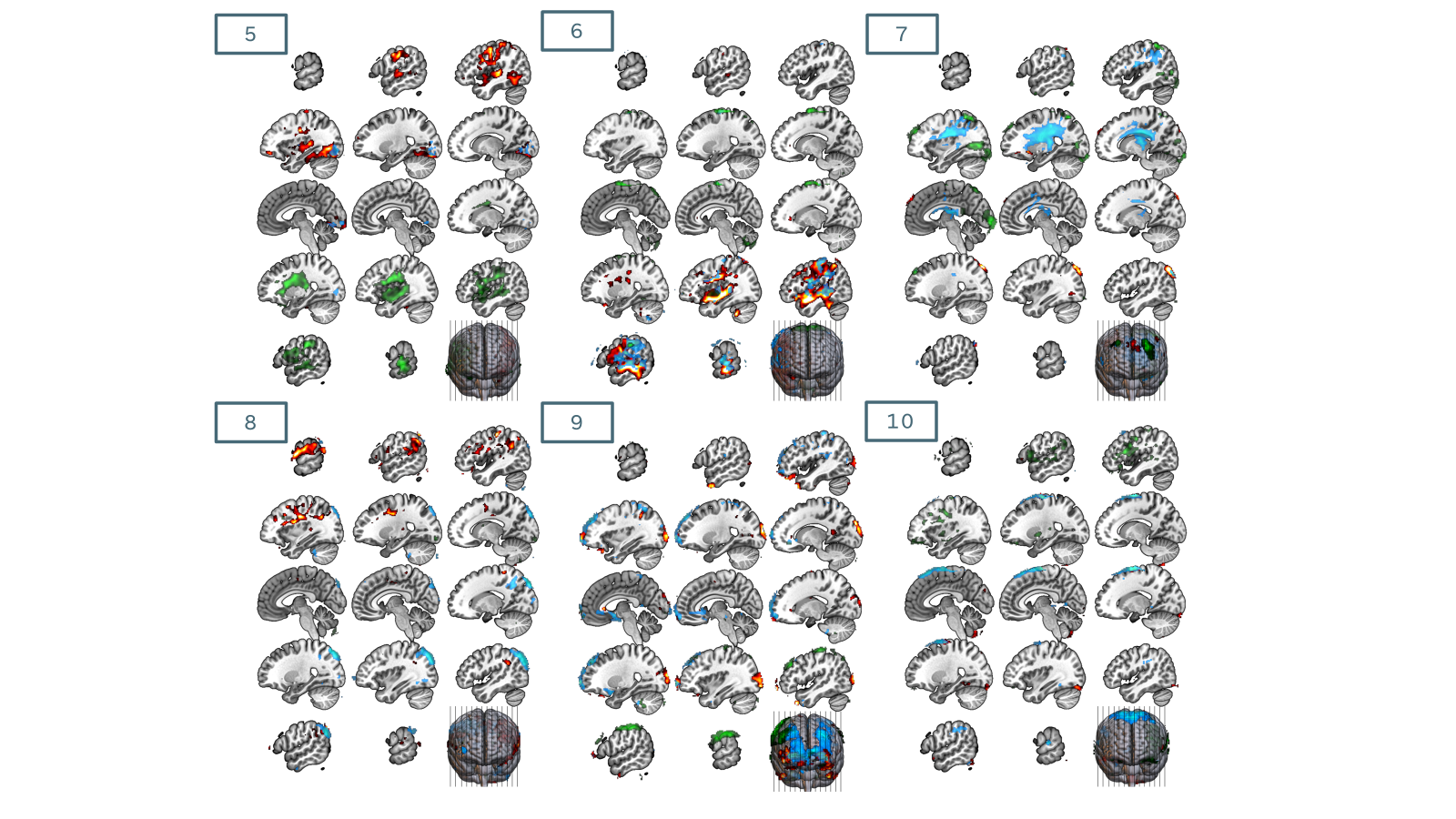

**Supplementary figure 2. Components 5-10 from ICA analysis.**
